# A novel screening method using CRISPRa and FM 1-43 to identify cation channels

**DOI:** 10.64898/2026.07.02.736146

**Authors:** RA Pak, NW Villarino, KL Hung, Y Wang, A Patapoutian

## Abstract

The discovery of sensory ion channels, such as thermosensitive transient receptor potential (TRP) channels and mechanosensitive PIEZOs, have transformed our understanding of mammalian sensory biology. However, the sensory receptor landscape remains incomplete, as many physiologically relevant sensory stimuli still lack identified molecular targets. Here, we describe a novel screening strategy utilizing FM 1-43, a fluorescent marker for activity of various cation channels, with a CRISPRa library (MPCL) targeting multi-transmembrane domain proteins. We validate this method by focusing on allyl isothiocyanate (AITC) and its putative receptor TRPA1. Specifically, we show that CRISPRa-mediated overexpression of TRPA1 is sufficient for FM 1-43 labeling when co-treated with AITC. Furthermore, we show that using FM 1-43 and AITC, we can efficiently FACS enrich TRPA1-expressing cells from a pool of MPCL-expressing cells. Collectively, this presents a novel method for rapidly screening select cation-dependent sensory stimuli.

## Introduction

The identification of sensory ion channels, such as thermosensitive TRP channels^1–5^ and mechanosensitive PIEZOs^6^, have been impactful in understanding mammalian sensory biology. Historically, discovery of these sensory channels has relied on a multitude of screening approaches, including cDNA expression libraries derived from dorsal root ganglion neurons1, bioinformatics to identify homologous proteins of known TRP channels2,4, and siRNA screens to assay mechanosensitive cell lines^6–8^. While such strategies have been historically successful, their efficacy for future studies faces major hurdles.

Large cDNA libraries stemming from transcriptomics or bioinformatics are cumbersome to maintain sequence fidelity, and make assumptions that they faithfully capture relevant mRNAA sequences and gene candidates. Functional assays like electrophysiology are labor intensive and limit throughput, while calcium imaging requires real-time recording to capture phenotypes. Critically, reverse genetic approaches to identify novel sensory ion channels, such as those used in Neuro2a cells for PIEZOs, are limited by the scarcity of culturable cell lines that exhibit stimulus-dependent currents. Considering the multitude of uncharacterized membrane proteins, some of which may encode novel cation-permeable proteins, and abundance of relevant physical, temperature, and chemical stimuli without a putative sensor or receptor, a need to develop a novel method to accelerate screening remains. Pooled genetic screens have been an effective and popular strategy^9–11^ to identify molecular targets for small molecules^12,13^ and sensory compounds^1^. Building on this, we sought to develop a broadly-applicable screening library to interrogate membrane proteins as potential receptors of small molecules, including chemosensory compounds and orphan drugs.

Over the past decade, CRISPR/Cas9-based methods have transformed our ability to manipulate and engineer cells^14^. In particular, genetic overexpression by CRISPR activation (CRISPRa)^15^, which drives endogenous gene expression via nuclease-dead Cas9 and transcriptional activators, has enabled a variety of methods to study protein targets and interactions, including pooled screening methods with fluorescent reporters^16^. The efficiency of pooled genetic screens has enabled a range of studies difficult to achieve with arrayed screens, including genome-wide studies^17^. Given the efficacy of CRISPRa for high-throughput screening, we sought to develop a novel pooled library of membrane proteins (MultiPass CRISPRa Library, MPCL) to identify novel ion channel targets of various stimuli. Furthermore, to stably translate transient ion channel activity into a permanent fluorescent marker, we combined the expression of this library with a fluorescent marker of cation channel activity, FM 1-43^18^.

## Results

FM 1-43 is a marker for non-selective cation channel activity in cultured cells^18,19^, known to be permeable to a diversity of ion channels, including various TRP channels, PIEZOs^19,20^, P2X receptors^18,21^, and TMC1/TMC2^18,22^. Cells labeled by FM 1-43 are stably fluorescent for several hours, allowing them to be isolated via fluorescence activated cell sorting (FACS), eliminating the kinetic restraints of real-time assays like electrophysiology and calcium imaging. Leveraging this property, we couple CRISPRa-driven over-expression of candidate membrane proteins with FM 1-43 labelling under defined stimuli. Cells which express ion channel candidates that respond to a given stimulus will be labeled by FM 1-43. FACS isolated cells can then be deep sequenced to identify enriched sgRNAs.

To demonstrate the feasibility of this approach we assessed FM 1-43 uptake with CRISPRa-mediated overexpression of TRPA1^3^, a well-characterized TRP channel. Using the hTERT-RPE-1 cell line, an immortalized human retinal pigmented epithelial cell line previously engineered to express the SunTag CRISPRa system (referred to as RPE-1 CRISPRa)^13^, we lentivirally targeted the human *TRPA1* gene, or the yeast-specific gene Gal4 as negative control. Following the previously described FM 1-43 fluorescence assay^19^, TRPA1-expressing cells co-treated with FM 1-43 and 100 uM AITC, a known chemical agonist of TRPA1, showed 6-fold increase in FM 1-43 fluorescent signal compared to control (Figure 1A-B), demonstrating that CRISPRa-mediated gene expression is sufficient to enable previously demonstrated TRPA1-dependant FM 1-43 labeling^23,24^.

**Figure 1:**
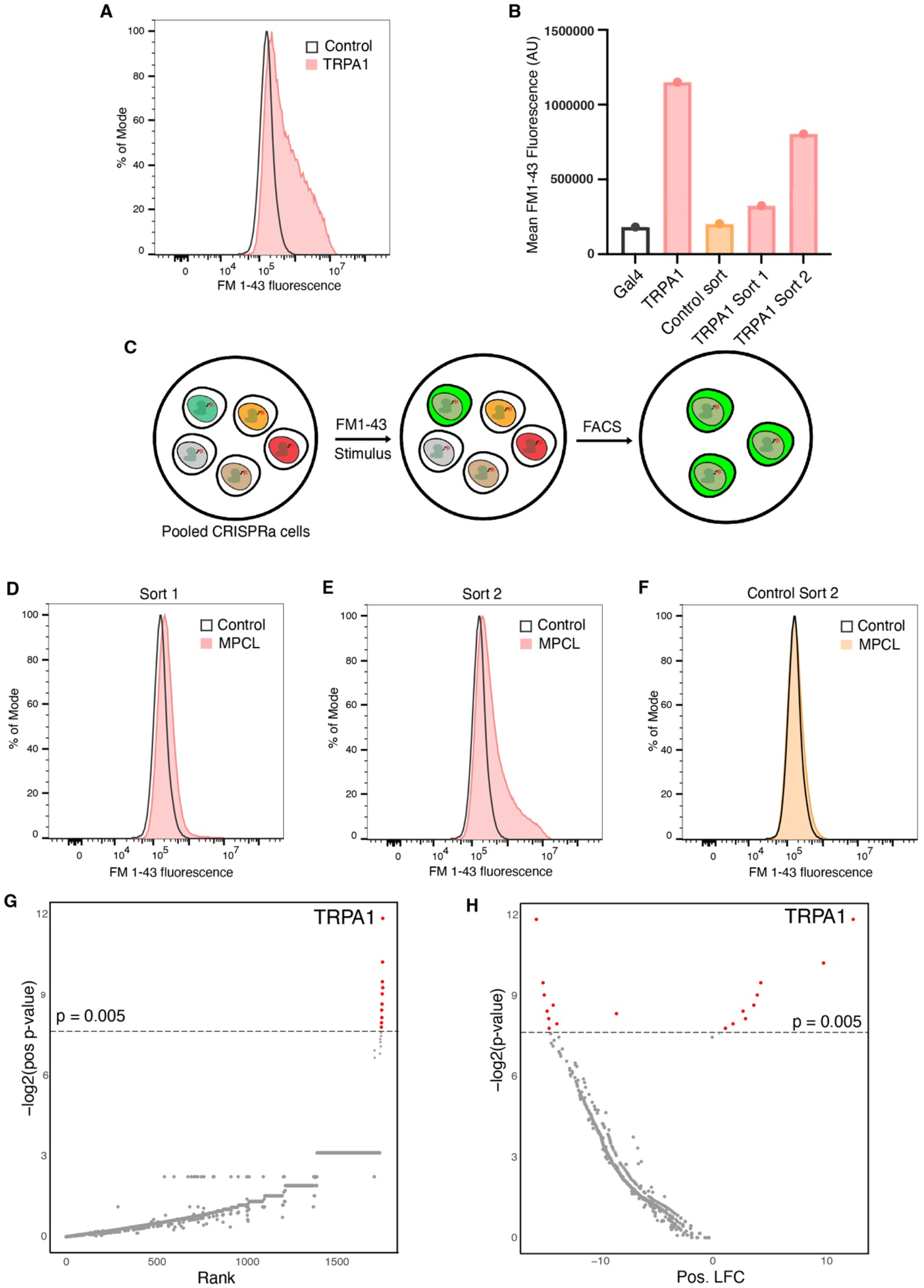
(A) Representative flow cytometry plots from RPE-1 CRISPRa cells expressing TRPA1 (red) or control (black line) treated with AITC (B) Quantification of mean FM 1-43 fluorescence (arbitrary units) from flow cytometry for tested conditions (C) Schematic of pooled CRISPRa screening. Pooled CRISPRa cells (left) are treated with FM 1-43 and stimulus, labeling only cells expressing the relevant sensor (middle). Labeled cells are FACS isolated (right) (D) Representative flow cytometry plots from MPCL RPE-1 CRISPRa cells after 1 round of FACS enrichment with AITC treatment (red), compared to control (black line) (E) Representative flow cytometry plots from MPCL RPE-1 CRISPRa cells after 2 rounds of FACS enrichment with AITC treatment (red), compared to control (black line) (F) Representative flow cytometry plots from MPCL RPE-1 CRISPRa cells after 2 rounds of FACS enrichment with mock treatment (orange), compared to control (black line) (G) sgRNA enrichment plot for samples shown in (E), with genes plotted by enrichment rank order (x-axis) with associated p-value (y-axis) (H) Data shown in (G), plotted as log fold chance (LFC, x-axis) with associated p-value (y-axis)

Given these results, we sought to leverage the long-term labeling properties of FM 1-43 with an entire library of pooled CRISPRa sgRNA expressed in RPE-1 cells. We first generated a list of human membrane proteins known or predicted to have multiple transmembrane domains (TMD), as virtually all ion channels have 2 or more TMDs, which yielded a total of 2933 targeted genes (Supplemental File 1). Three sgRNA targeting each gene were cloned into dual sgRNA vectors^25^, and transduced into RPE-1 CRISPRa cells.

Considering CRISPRa-mediated TRPA1 expression was sufficient for FM 1-43 labeling with AITC, we reasoned that TRPA1 could be enriched or isolated from the MPCL RPE-1 CRISPRa pool of cells by FACS with chemical agonism and serve as a proof-of-concept (Figure 1C). We treated MPCL-transduced cells with FM 1-43 and AITC, which did not initially reveal a distinct FM 1-43-positive population (Figure 1D), consistent with TRPA1’s expected low representation of around 1 in 3000 cells. Using FACS, we isolated the brightest 2% of cells (referred to as “Sort 1”). These cells were expanded in culture and re-treated with FM 1-43 and AITC, demonstrating a 1.78-fold increase in FM 1-43 fluorescence compared to control. We sequentially enriched these cells by again FACS-isolating and culturing the brightest 2% of cells (Referred to as “Sort 2”), which then demonstrated 4.4-fold increase in FM 1-43 fluorescence with AITC compared to control. In contrast, DMSO-treated controls showed negligible changes to FM 1-43 fluorescence after two rounds of equivalent sorting (Figure 1B, F).

Next, to deconvolute the enriched populations and verify that this AITC-dependent FM 1-43 labeling was through TRPA1, we subjected AITC-enriched and negative control cell populations to sgRNA deep sequencing and analyzed relative sgRNA abundance. We observed some overlap in enriched genes between experimental and control conditions, perhaps due to FM 1-43’s sensitivity to changes in vesicular membrane flux (Supplemental File 3). However, by calculating the fold-change of sgRNA enriched in AITC-treated cells compared to sgRNA enriched in mock-treated cells, we effectively filtered out background enrichment. Our analysis identified 11 genes specifically enriched in the AITC-treated cells, where *TRPA1* emerged with the highest fold-enrichment (Figure 1G-H, Supplemental File 3). While the other 10 candidates were not further validated, they can largely be attributed as background or false positives, as TRPA1 is known to be the putative receptor for AITC in sensory neurons. Collectively, these results demonstrate that this pooled screening approach with FM 1-43 is useful for identification of protein targets of sensory stimuli.

## Discussion

This study establishes a rapid, adaptable screening strategy that merges FM 1-43–based fluorescence sorting with pooled CRISPRa libraries to identify cation-permeable sensory channels. By harnessing the efficiency of CRISPRa while bypassing the kinetic constraints of electrophysiology and calcium imaging assays, our approach accelerates the initial discovery phase. Although downstream electrophysiological validation remains essential, this platform may significantly reduce the screening bottleneck.

Notably, this method in its current form is limited by single genetic perturbations, and cannot capture novel sensors with multiple distinct subunits, unless endogenously expressed in that cell. Considering the efficacy of multiple gene perturbations in other works^16,26^, future iterations of MPCL may similarly perturb 2 or more genes simultaneously. Furthermore, FM 1-43 is thought to label cells expressing a subset of non-selective cation channels through an unknown mechanism^19^. As such, it remains beneficial to first test whether FM 1-43 can label cells expressing an endogenous current before attempting to identify the unknown ion channel in question. Nonetheless, given the calcium-dependent responses observed in DRG neurons for unresolved sensory stimuli, this novel method presents a high-throughput alternative for novel discovery.

Notably, our method generated a high false discovery rate (FDR) (Supplemental File 3), likely attributed to limited numbers of sgRNA targeting each gene. However, future enhancements— such as expanding sgRNA coverage or integrating orthogonal reporters—could bolster screen fidelity and analysis. Additionally, our sequencing analysis revealed that a number of genes are ubiquitously enriched with FM 1-43 treatment, irrespective of experimental parameters. Many of these genes are known to regulate or affect membrane composition and activity. Considering the amphipathic properties of FM 1-43, we reason that any genes affecting cell membranes may also affect FM 1-43 labeling. Indeed, most of the genes also enriched in the TRPA1 screen are membrane affecters.

We anticipate that this method will facilitate the discovery of novel ion channels and deepen our understanding of membrane protein function in various physiological contexts. Furthermore, MPCL’s design is broadly applicable to membrane protein research beyond sensory biology, including target identification for small molecules and mapping cell-surface interactions.

## Methods and Materials

### sgRNA Library cloning, deep sequencing preparation, and analysis

To generate a gene-list targeting membrane proteins with multiple TMDs, we mined solved and predicted structure data from UniProt, yielding a candidate list of 2933 human genes (circa 2019).

The top 3 ranked sgRNA targeting these genes from the hCRISPRa-V2 library^17^ were cloned into dual sgRNA vectors as previously described^25^, using sgRNA 1 + 2 and 1 + 3 as pairs, for a total of 5866 vectors. Additionally, 1000 non-targeting sgRNA vectors from hCRISPRa-V2 were cloned to serve as internal negative control, representing ∼15% of the total library. We refer to this library as the Multi-Pass CRISPRa Library (MPCL) (Supplemental File 2). Similar to previous experiments, we transduced the MPCL library into RPE-1 CRISPRa cells at 1000x coverage to ensure full library diversity, and a multiplicity of infection (MOI) of 0.3 to ensure any given cell is transduced no more than once. Transduced cells were FACS purified, yielding a final population of 90% purity.

Libraries for sgRNA sequencing were prepared following published protocols. Briefly, ∼5 million cells were collected from each sample. Total genomic DNA was purified using a NucleoSpin Blood Mini kit (Machery Nagel). Genomic DNA was directly amplified using a Nested PCR to first amplify sgRNA sequences using universal primers, then to add indices and hybridization sequences for sequencing (TruSeq). Amplicons were purified with AMPureXP beads (Beckman Coulter) and quantified with a Qubit 4 Fluorometer (Thermo Fisher). Libraries were sequenced on a NovaSeq (Illumina).

Read2 fastq files from paired-end sequencing containing guide sequence fragments were used for downstream analyses. Relative abundances of guide RNAs were measured using MAGeCK (v.0.5.9.4). Guide RNA counts were obtained using the “mageck count” command. Pairwise comparisons between treatment and control conditions were performed using the “mageck test-- paired” command. P values and false discovery rates from these pairwise comparisons were used for identifying enriched gene hits.

### Cell culture

hTERT-RPE-1 CRISPRa cells were previously engineered to express CRISPRa SunTag machinery, and were used for all CRISPRa experiments. Cells were cultured in DMEM/F-12 media, supplemented with 10% FBS and 100 U/mL Pen/Strep/Glutamine (Thermo Fisher), at 37C and 5% CO2. For MPCL-expressing cells, cultures were maintained at 1000x coverage of the library.

HEK293T cells (ATCC) were used for lentivirus production. Cells were cultured in DMEM (4.5 g/mL glucose) supplemented with 10% FBS and 100 U/mL Pen/Strep/Glutamine (Thermo Fisher), at 37C and 5% CO2.

### Lentivirus production and transduction

Lentivirus for sgRNA vectors were produced using a modified 3rd generation system. Briefly, HEK293T cells were transfected at 60-70% confluency in 6-well or 15 cm round dishes with lentivirus plasmids using Mirus (Millipore Sigma) or Lipofectamine 2000 (Thermo Fisher). Media was exchanged 12-16 hours later. Supernatant was collected, filtered, and frozen (-80C) 48 hours post-transfection.

RPE-1 cells were plated the day before lentivirus transduction to achieve 30-40% confluency. Lentivirus was thawed at room temperature until thawed, and mixed into culture supernatant with Polybrene (6 ug/mL) to enhance transduction. 12-16 hours later, media was exchanged. 72 hours post-transduction, cells were analyzed for BFP signal on an ACEA NovoCyte flow cytometer. Parameters were optimized to achieve 1000x coverage of the full MPCL library, and an MOI of 0.3 to ensure single transduction events per cell.

### FM 1-43 fluorescence assay

We used the FM 1-43 fluorescence assay as previously described^11^. Briefly, 10 uM FM 1-43FX (Invitrogen), 500 uM Advasep-7, and 100 uM AITC were resuspended in Hank’s Buffered Saline Solution (HBSS, 10 mM HEPES, pH 7.4). Cells were co-treated with FM 1-43 and AITC for 15 minutes at room temperature, then washed twice with Advasep-7 on an orbital shaker for 5 minutes each. For mock treatment, equivalent volumes of DMSO were used in place of AITC. Cells were disassociated using standard Trypsin protocols (Thermo Fisher) and re-suspended in FACS buffer (PBS supplemented with 2% FBS and 1 mM EDTA) in polystyrene tubes. Cells were analyzed on an ACEA NovoCyte, or sorted on a MoFlo Astrios EQ (Beckman Coulter) for mean FITC-A (488 nm excitation, 525 nm emission) fluorescence as readout of FM 1-43 labeling. Resulting data was analyzed in FlowJo. Sorted cells were cultured and expanded for up to two weeks between enrichment rounds to achieve 1000x coverage of the sorted population.

## Acknowledgements

RAP and AP were responsible for the conception, design, and interpretation of the experiments. NWV assisted with experimental design and execution. KLH analyzed deep sequencing results. YW analyzed UniProt data for transmembrane domain counts.

The authors thank Marco Jost for expertise and many helpful discussions. The authors additionally thank the Flow Cytometry Core at the Scripps Research Institute for their expertise and assistance; the Genomics Core at the Scripps Research Institute; and the IGM Genomics Center at University of California, San Diego.

## Notes

### Competing Interest Statement

The authors have declared no competing interest.

